# New algorithms for accurate and efficient de-novo genome assembly from long DNA sequencing reads

**DOI:** 10.1101/2022.08.30.505891

**Authors:** Laura Gonzalez-Garcia, David Guevara-Barrientos, Daniela Lozano-Arce, Juanita Gil, Jorge Díaz-Riaño, Erick Duarte, Germán Andrade, Juan Camilo Bojacá, Maria Camila Hoyos, Christian Chavarro, Natalia Guayazan, Luis Alberto Chica, Maria Camila Buitrago Acosta, Edwin Bautista, Miller Trujillo, Jorge Duitama

## Abstract

Producing de-novo genome assemblies for complex genomes is possible thanks to long-read DNA sequencing technologies. However, maximizing the quality of assemblies based on long reads is a challenging task that requires the development of specialized data analysis techniques. In this paper, we present new algorithms for assembling long-DNA sequencing reads from haploid and diploid organisms. The assembly algorithm builds an undirected graph with two vertices for each read based on minimizers selected by a hash function derived from the k-mers distribution. Statistics collected during the graph construction are used as features to build layout paths by selecting edges, ranked by a likelihood function that is calculated from the inferred distributions of features on a subset of safe edges. For diploid samples, we integrated a reimplementation of the ReFHap algorithm to perform molecular phasing. The phasing procedure is used to remove edges connecting reads assigned to different haplotypes and to obtain a phased assembly by running the layout algorithm on the filtered graph. We ran the implemented algorithms on PacBio HiFi and Nanopore sequencing data taken from bacteria, yeast, *Drosophila*, rice, maize, and human samples. Our algorithms showed competitive efficiency and contiguity of assemblies, as well as superior accuracy in some cases, as compared to other currently used software. We expect that this new development will be useful for researchers building genome assemblies for different species.

## INTRODUCTION

Contiguous and accurate assembly of complex eukaryotic genomes is one of the most challenging tasks in current biotechnology and bioinformatics (Baker M 2012, Nurk et al. 2022). Bioinformatic tools for genome assembly are used to sort and orient partial reads produced by various sequencing technologies. Partial genome assemblies, including most gene-rich regions, have been generated in the last decade. However contiguous and high-quality assemblies are required to integrate synteny information in genome-scale comparative genomics and pangenomics, to study evolution and dynamics of mobile elements, for population genomic analysis, such as genome-wide association studies (GWAS), and for the discovery of genomic footprints of selection (Amiri et al. 2018, Xu et al. 2020). High-quality assemblies are also useful to understand the genome evolution of species (Hu et al. 2021), to identify structural variations (Ouzhuluobu et al. 2020), and to define the gene repertoire including targets for resistance in plants and animals, as well as virulence factors and effectors in pathogens (Bhadauria et al. 2019). This complete gene catalog is key for identifying interesting genomic target regions for plant and animal breeding (Low et al. 2020, Song et al. 2021), and personalized medicine. Moreover, genome assemblies have been useful in pathogen surveillance for public health (Taylor et al. 2019).

The production of sequencing data has grown exponentially in the last years and genome assembly has become a routinary task; however most of the currently available genomes have been sequenced using high-quality short-read technologies such as Illumina. Currently, long-read technologies, such as PacBio and Nanopore, have improved the quality of data and allowed a better de novo assembly of genomes, haplotype phasing, and structural variants identification (Hon et al. 2020). Nanopore sequencing technologies offer the advantage of producing the longest read lengths (Mbp range), the more common lengths being 10 to 30 Kb, as these are limited by the quality and size of the DNA delivered to the sequencing pore (Amarasinghe et al. 2020). Furthermore, some of the Nanopore sequencers can be portable and generate data in real time, proving useful for field research and diagnostics (Xu and Seki 2019). In contrast, PacBio single molecule real-time (SMRT) sequencing delivers reads of 30 Kb on average, it has a low coverage bias across different values of G+C content, and allows for the direct detection of DNA base modifications (Nakano et al. 2017). Nanopore and PacBio CLR long-reads have an average error rate of ∼15%. Nevertheless, PacBio has developed a new method to generate HiFi reads (high-fidelity reads with an average error rate of ∼0.5%), which allow the assembly of complete chromosomes, even for diploid or polyploid organisms. Despite the high assembly contiguity achieved with long reads, other strategies can be used to improve assemblies, such as: Hi-C (Zhou et al. 2019), parental information (Wenger et al. 2019), and Strand-seq (Hills et al. 2021).

Most of the commonly used tools to assemble long-read datasets implement the overlap-layout-consensus (OLC) algorithm. These were developed to assemble reads with high error rates, such as the Nanopore and PacBio CLR reads. Canu (Koren et al. 2017) uses a MinHash overlapping strategy (Berlin et al. 2015) with a tf-idf weighting to identify overlaps. Then, a linear graph is constructed using a greedy best-overlap algorithm. WTDBG (Ruan and Li 2019) implements minimizers for efficient identification of overlaps. Flye (Kolmogorov et al. 2019) implements an algorithm to resolve repeats from a possibly inaccurate initial assembly. FALCON (Chin et al. 2016) implements a simple haplotype phasing algorithm to perform read clustering and to generate phased assemblies. After the emergence of PacBio HiFi reads, new algorithms have been developed to perform error correction. These algorithms aim for perfect reads in which single nucleotide differences can be used to resolve differences between repetitive elements (Nurk et al. 2020, Cheng et al. 2021). HiCanu is an improvement of Canu that implements homopolymer compression to align and correct reads having base counts on homopolymer tracts as main source of error (Nurk et al. 2020). HiFiASM integrates haplotype phasing to perform haplotype aware error correction (Cheng et al. 2021). Error correction of long reads, especially Nanopore reads, remains an important step during genome assembly and is usually a computationally expensive process. NECAT was developed as an error corrector and de novo assembler for Nanopore reads (Chen et al. 2021). In NECAT error correction is based on a two-step progressive method by which low-error-rate subsequences of reads are corrected first, and then they are used to correct high-error-rate subsequences.

In this work we introduce a new software implementation for genome assembly from long-read sequencing data. It includes new algorithmic approaches to build overlap-layout-consensus (OLC) assembly graphs, and to identify layout paths. Benchmark experiments on PacBio HiFi and Nanopore data from organisms of different species including *Escherichia coli*, yeast, *Drosophila melanogaster*, rice, maize, and human show that our algorithms are competitive and, in some cases, more accurate, compared to previous solutions. These algorithms are implemented as part of the Next Generation Sequencing Experience Platform (NGSEP) (Tello et al. 2019), allowing a tight integration with genome comparison and detection of genomic variants within a single easy-to-use tool for analysis of both short and long read DNA sequencing data.

## RESULTS

### K-mer count based hashing for efficient and accurate construction of assembly graphs

We implemented a new hashing scheme for minimizers to efficiently identify overlaps and build OLC graphs. Figure 1 shows the implemented algorithm to build an overlap graph and a layout. The graph construction is similar to that of the Best Overlap Graph (Miller et al. 2008), having two vertices for each read representing the start (5’-end) and the end (3’-end) of the read. In this representation, the graph does not need to be a multigraph. Let X^s^ and X^e^ be the two vertices generated from each read X. If the end of read A has an overlap with the start of read B, this overlap is represented with the edge {A^e^,B^s^}. Conversely, if the end of read A has an overlap with the start of the reverse complement of B, this overlap will be represented by the edge {A^e^,B^e^}. In our representation, the graph is completely undirected to take into account that reads are sequenced from the two strands of the initial template with equal probability and hence, there is no a-priori information on which one should be considered the positive strand.

**Figure 1.**
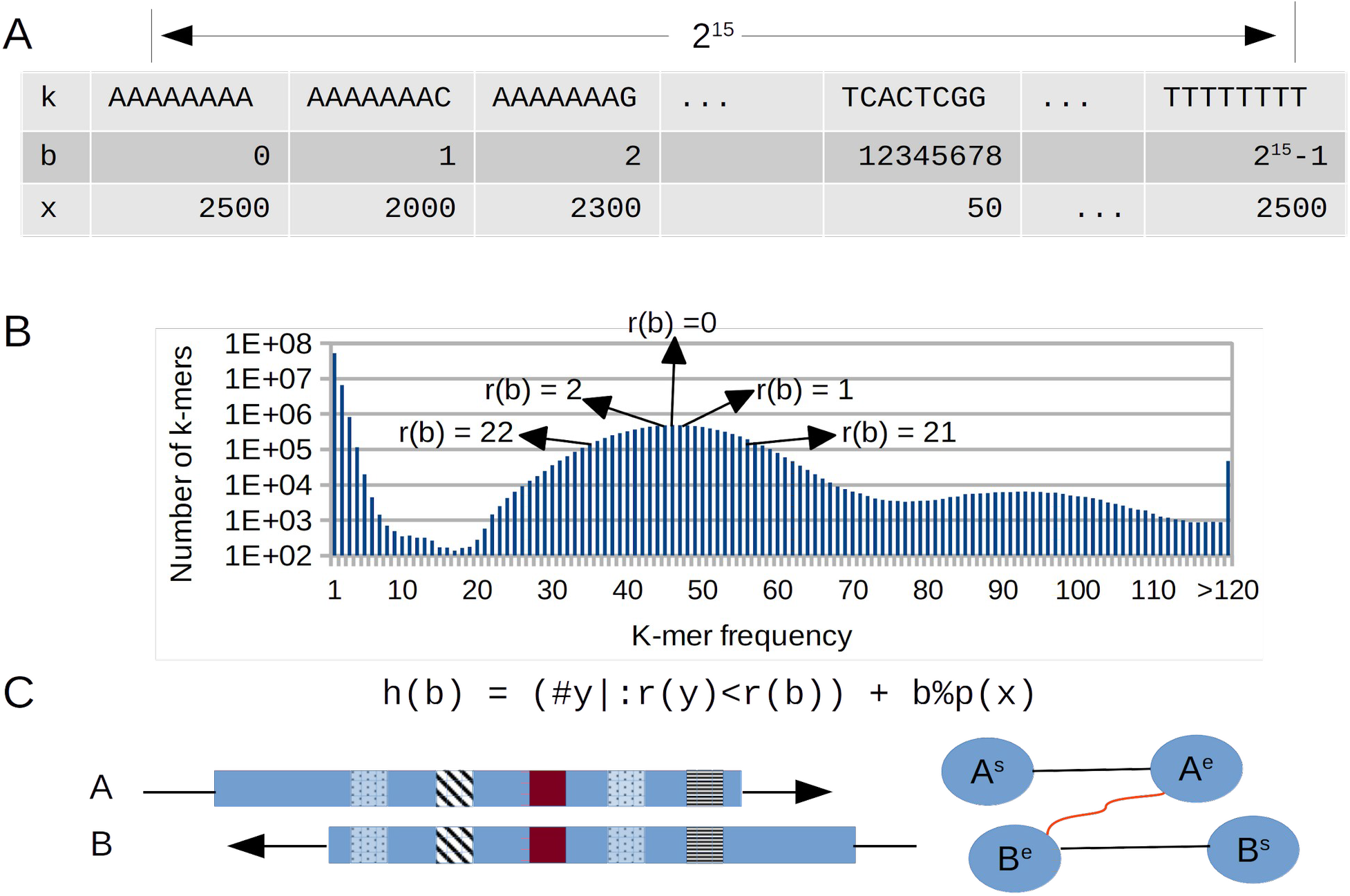
Overview of the graph construction algorithm implemented in NGSEP for de-novo assembly of long reads. A. Fixed array to calculate counts of 15-mers. B. The distribution of k-mer frequencies is used to rank edges based on their distance from the peak corresponding to single copy regions. C. A hash value is calculated from the rank to select minimizers and identify overlaps. Dynamic programming is used to cluster k-mer hits.

Similarly to the graph construction implemented in WTDBG (Ruan and Li 2019), we built a minimizers table from the reads, to identify overlaps in linear time relative to the total number of sequenced base pairs. However, we implemented a different procedure to calculate hash codes, that changes the priority to select k-mers as minimizers. Before calculating minimizers, we first build a 15-mer spectrum table, calculating the count distribution across the reads. Analyzing this distribution, the algorithm infers the mode that corresponds to the average read depth, and estimates the assembly size. To achieve an efficient calculation of the k-mer distribution, the spectrum table is built with a fixed k-mer length of 15 (instead of the input k-mer length used later), because that is the maximum length to create the table as a fixed array of length 2^30^ in which the index of the array corresponds to a unique encoding of each possible DNA k-mer. The data type of this array is a two-byte integer to store a count per k-mer up to 2^15^, which is enough for real whole genome sequencing datasets. This implementation ensures a fixed memory usage of 2^31^ bytes (about 2 gigabytes), regardless of the input size and genome complexity. The 15-mer spectrum allows not only to approximate the assembly length and average read depth, but also to calculate the hash value of read k-mers.

To identify overlaps, k-mers of a user-defined length (up to 31) are calculated for each read. Each k-mer is uniquely encoded as a 62-bit number *b* and the count *x* of the 15-mer suffix on the 15-mer spectrum is calculated. A rank r(*x*) is calculated from the count, as two times the distance from the mode corresponding to the haploid number. The hash value h(*b*) is calculated as the number of k-mers with rankings smaller than r(*x*) plus the module of the division between *b* and the smallest prime number larger than *x*. This last term is a simple scheme to simulate randomness for k-mers within the same rank. This hashing scheme allows the prioritization of real k-mers that are likely to come from single-copy regions of the haploid genome during the calculation of minimizers. At the same time, k-mers from repetitive regions have larger hash codes, which reduces their priority to become minimizers but does not discard them completely. We implemented a simulated alignment of each candidate overlap to calculate different measures associated with each edge in the overlap graph, avoiding a complete pairwise alignment between candidate pairs at this stage of the process. First, matching k-mers (minimizers) between a subject (longer) read and a query (shorter) read are clustered based on consistency of the prediction of overlap start that can be inferred from the relative location of the k-mer in the subject sequence. Assuming that indel errors are randomly distributed across the two sequences and that insertion and deletion errors have a similar probability of occurrence, the inferred starting point for k-mers corresponding to a real overlap should be consistent (have a low variance). Conversely, inferred starting points for matching k-mers supporting false positive overlaps due to repetitive structures (up to a certain length) should have a larger variance. We implemented a clustering procedure similar to k-means to group k-mer hits that are likely to support the same alignment, using the inferred starting points as centroids. The average number of k-mer hits for each k-mer is used to infer the number of different clusters that can be expected. Up to two clusters with the largest k-mer count are retained as long as they support two of the four possible alignment configurations (start-start, start-end, end-start, and end-end). Because an overlap length can also be inferred from each matching k-mer, the overlap for a cluster of matching k-mers is inferred as the average of the inferences performed from each matching k-mer.

### Layout construction as an edge selection problem

The statistics collected during the simulated alignment step are used during the layout stage to select edges that will be part of the assembly paths. For each edge, derived from a k-mers cluster, relevant statistics include the predicted overlap, the number of shared k-mers building the overlap, the number of base pairs from the subject sequence covered by the shared k-mers (CSK), and the first and the last position of both the subject and the query sequence having k-mers supporting the possible overlap. The layout algorithm ranks and selects edges based on the knowledge that can be inferred from the distribution of the different statistics. Although in a real experiment true layout edges are unknown, we first identify edges that are reciprocal best for their corresponding vertices, both in terms of overlap length and CSK, and that connect vertices with total degree less than three standard deviations from the average. These edges are termed “safe” and it is assumed that they will be part of the layout. Because they are reciprocal best, these edges will generate an initial series of paths within the graph. Moreover, it is assumed that the distribution of overlap length and CSK calculated from these edges would be a good representation of the distributions calculated from all true layout edges. The cost of each remaining edge is calculated as a likelihood of the edge features given the distributions inferred from the safe edges. Whereas a normal distribution is fitted for the overlap and the CSK, a beta distribution is fitted for the proportion of overlap calculated from the first and the last overlap position supported by k-mers. Likelihoods are calculated as p-values of the edge features. Log-likelihoods of the features are added to calculate the total edge likelihood and sort edges based on this feature. Edges are then traversed in descending order to augment the paths initially derived from safe edges. An edge is selected if it does not include an internal path vertex and if it does not create a cycle. Figure 2 shows a schematic diagram of this procedure.

**Figure 2.**
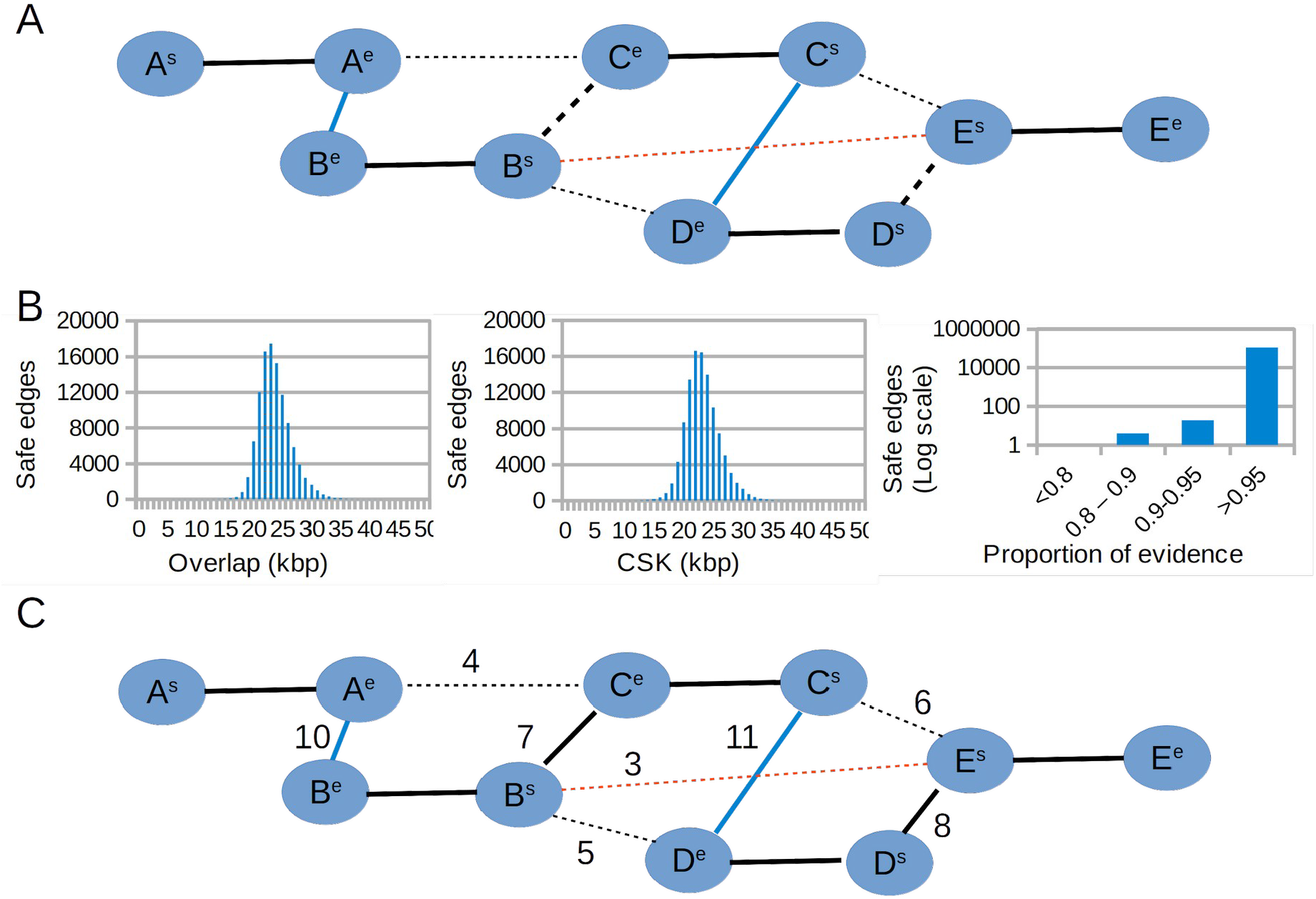
Layout algorithm. A. Safe edges (blue) are selected as reciprocal best in both overlap and coverage of shared Kmers (CSK). The red edge represents a false positive. Bold solid black edges connect vertices of the same read. Bold dashed edges are true layout edges that are not reciprocal best. Other dashed lines represent true non-layout edges. B. Distributions of overlap, CSK and proportion of evidence for safe edges of the rice 20 Kbp PacBio HiFi data (details in the next section). C. Log likelihoods are calculated for each edge based on the distributions; layout edges not selected in the first step are selected based on their ranking.

Once paths are constructed, an initial consensus is built concatenating layout vertices. On each step, the next read is aligned to the consensus end to recalculate the true overlap and the consensus is augmented with the substring corresponding to the overhang of the alignment. At the same time, embedded reads are recovered and mapped to the consensus. Once all reads are mapped, the following polishing algorithm is executed to improve the per base quality of the assembly: first, pileups are calculated for each position to identify the base with the largest count and update the consensus if needed. Then, similar to the process to call variants, a second step calculates “active regions” across the alignment, which are defined as contiguous regions in which each base pair is at most 5 bp away from an indel call. Once active regions are calculated, a de-Bruijn graph is built from the read segments spanning the active region and a mini-assembly is executed to calculate the corrected segment.

### Benchmark with PacBio HiFi data

To test the performance of NGSEP with PacBio HiFi data, we assembled genomes from publicly available HiFi reads of the indica rice variety Minghui 63 (15 Kbp and 20 Kbp reads), the B73 maize inbred line, and the human cell line CHM 13 using NGSEP and three commonly used tools (Canu, Flye, and HiFiASM). Figure 3 shows the results of these benchmark experiments. The contiguity of each assembly, measured as the Nx curve, is contrasted with the number of misassemblies against a curated reference genome, as measured by Quast (Gurevich et al. 2013). The complete statistics are available in the Supplementary Table T1. Regarding the rice data, the assemblies generated by HiFiASM and NGSEP have the highest N50 values for the 15 Kbp and 20 Kbp datasets respectively. In both cases, at least 95% of the genome (395 Mbp) was assembled in less than 20 contigs. Canu ranks third, close to NGSEP for the 15 Kbp dataset and close to HiFiAsm for the 20 Kbp data. Flye shows the lowest contiguity in all datasets (Figure 3A). Conversely, for the maize and the CHM13 datasets, the assemblies generated by NGSEP have lower contiguity compared to those generated by HiFiAsm and Canu, but still have better contiguity compared to the assemblies generated using Flye. For the maize dataset all the tools assembled the genome in more than 500 contigs with minimum length of 50 Kbp. Using this dataset, the N50 value ranged from 4.4 Mbp (Flye) to 37 Mbp (HiFiASM). This is probably caused by a lower average read depth and higher complexity, as compared to the rice datasets. The same behavior was observed in the human cell line where the assembled genomes were highly fragmented and the N50 value ranged from 29 Mbp (Flye) to 86 Mbp (HiFiASM).

**Figure 3.**
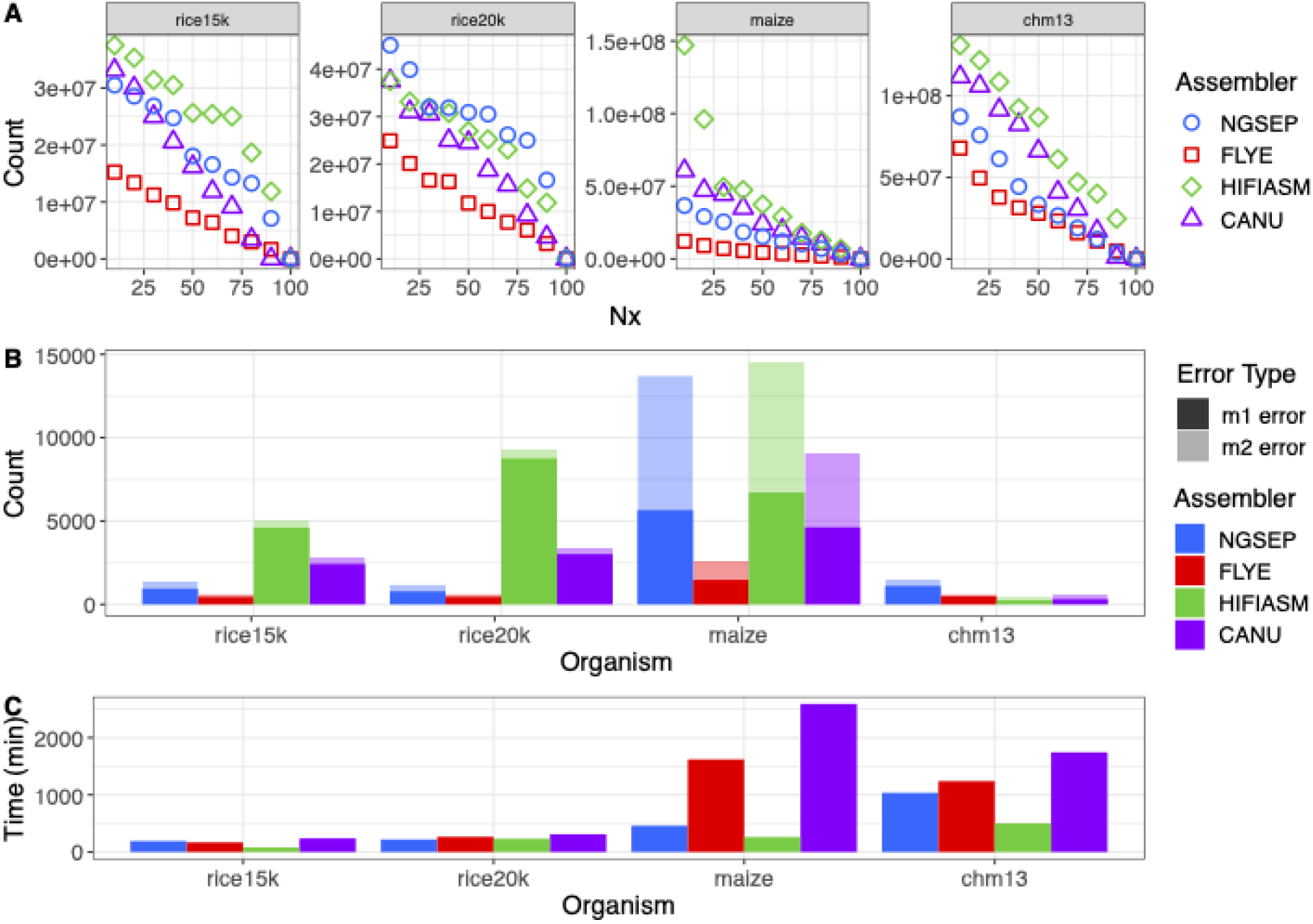
Assembly results for haploid or inbred samples. A. Nx curve B. Misassemblies(m1 error) and local misassemblies (m2 error) reported vs reference genomes. Rice15k corresponds to *Oryza sativa* 15k HiFi reads, rice20k to *O. sativa* 20k HiFi reads, maize corresponds to *Zea mays B73* HiFi reads, and chm13 corresponds to the human cell line chm13. C. Execution time (in minutes) for each experiment. Figure 3B shows the number of misassembly errors identified by Quast, using a curated reference genome for comparison. Errors are classified as long-range misassemblies (m1) and local misassemblies (m2). With the exception of the maize assemblies produced by NGSEP and HiFiAsm, most assemblies reported more m1 errors than m2 errors. Flye assemblies reported the lowest numbers of misassemblies for the plant samples, whereas the HiFiAsm assembly reported the lowest number for CHM13. Conversely, HifiASM assemblies reported the highest total number of misassemblies for plant samples. The number of errors in assemblies generated with NGSEP on the rice samples was about 1.6 times higher than the number of errors generated by Flye, but it was up to 5 times lower than the number of errors generated by HiFiAsm. Additionally, in the maize sample, NGSEP generated fewer misassemblies than HiFiAsm.

Regarding computational efficiency, Figure 3C shows a comparison of the runtimes (having available 32 threads) required by each tool to assemble each of the datasets. HiFiAsm and Canu are consistently the fastest and the slowest tools respectively. NGSEP requires a lower runtime than Flye in all datasets except for the rice 15 Kbp dataset, where Flye finishes 24 minutes faster than NGSEP. In absolute numbers, NGSEP is able to assemble the rice datasets in less than 4 hours, the maize dataset in less than 8 hours, and the CHM13 dataset in less than 18 hours.

Combining the evaluation of accuracy and efficiency, NGSEP has better computational efficiency than Flye and Canu and the assemblies have better contiguity than those of Flye, and fewer misassemblies than most of those assembled using Canu. Compared to HiFiAsm assemblies, NGSEP assemblies of plant samples have lower error rates and the 20 Kbp NGSEP assembly showed the best contiguity for rice.

### Assembly and haplotyping of diploid samples

We integrated our previous implementation of the ReFHap and the DGS algorithms to perform single individual haplotyping of diploid heterozygous samples (Duitama et al. 2012). Unlike the previous implementation, which received a non-standard file with base calls for each heterozygous site, the two algorithms can now be executed from the VCF file with individual genotype calls and a BAM file with long reads aligned to the reference genome and sorted by reference coordinates. Moreover, we integrated the ReFHap algorithm within the assembly process of diploid samples to obtain phased genome assemblies from HiFi reads. ReFHap is executed independently on reads aligned to an initial assembly, which is generated using the methods described above for haploid samples. The goal of this phase is to identify and break edges in the assembly graph connecting reads sequenced from different haplotypes. Large deletions and regions of homozygosity larger than the read length usually break each contig into haplotype blocks (Cheng et al. 2021). Read depth within each block and between block boundaries is calculated to break the contig in contiguous regions classified as true phased regions, large heterozygous deletions, or regions with high homozygosity. Edges connecting reads within true phased regions and assigned to different haplotype clusters are removed from the assembly graph.

To validate the accuracy of the complete process to assemble phased genomes, we first simulated two single chromosome diploid genomes. The first was constructed from two publicly available MHC alleles. The second was constructed from the copies of the rice chromosome 9 corresponding to the Nipponbare and the MH63 assemblies. A high heterozygosity rate is expected in both cases. We assembled simulated reads from both individuals using both NGSEP and HiFiAsm. For the MHC haplotypes, NGSEP was able to reconstruct the reference allele in two contigs of lengths 4.4 Mbp and 0.3 Mbp, and the alternative allele in three contigs of lengths 3.5 Mbp, 0.5 Mbp and 0.2 Mbp (Supplementary figure S1). No switch errors (changes between real alleles within a contig) were detected in this assembly. Conversely, three contigs assembled by HiFiAsm, with lengths of 4.4 Mbp, 0.8 Mbp and 1.6 Mbp, mapped to the alternative MHC allele and one contig of 2.8 Mbp mapped to the reference MHC allele. Hence, the alternative allele was overrepresented, having the two smaller contigs embedded within the largest contig. The largest contig was also larger than the original allele because the left 100 Kbp could not be mapped and the right 200 Kbp was duplicated. Conversely, the reference allele was sub represented. Figure 4 shows the reconstruction of the rice alleles by NGSEP and HiFiAsm. NGSEP assembled one large contig having three switch errors and five additional contigs covering the regions not covered by the first contig. HiFiAsm assembled most of the MH63 chromosome in two contigs and most of the Nipponbare chromosome also in two contigs. Three switch errors were detected in this case. It also produced four small contigs (about 200 Kbp), two of them overlapping with longer contigs.

**Figure 4.**
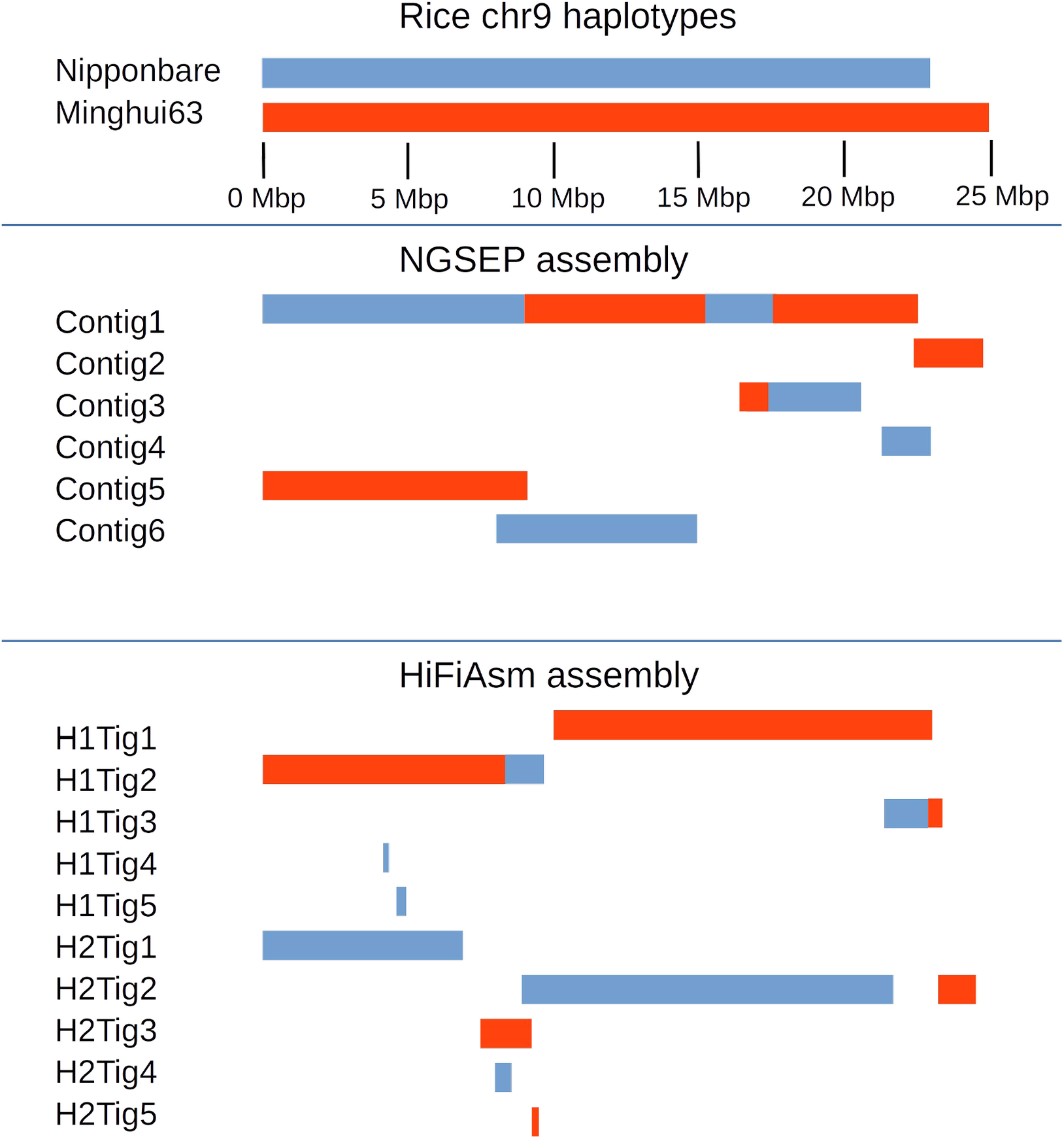
Results of a diploid assembly of a simulated diploid individual built from the chromosome 9 sequences of the rice japonica accession Nipponbare and the Indica accession Minghui63. Blue blocks show Nipponbare haplotypes, whereas red blocks indicate Minghui63 haplotypes. Changes in color in the same row represent switch errors.

To further assess the performance of NGSEP assembling diploid samples, we executed assemblies from publicly available HiFi reads of the human individual HG002. NGSEP generated an assembly with a total length of 5,593.63 Mbp distributed into 12,318 contigs. The NGA50 was 1.68 Mbp. In contrast, HiFiAsm produced an assembly of 5,979.17 Gbp distributed into 851 contigs and a NGA50 of 91.07 Mbp. Despite the large difference in contiguity, we also collected some of the metrics proposed by Cheng et al. 2021, related to the ability of the assembly to reconstruct the two alleles of each gene present in the diploid sample (Table 1). For the case of HiFiAsm, we calculated the metrics for both the primary assembly and the phased assembly. From the 35,547 single-copy genes in the reference genome, NGSEP recovered 81% of them, and HiFiASM recovered 89% in the phased assembly and 98% in the primary assembly. However, the NGSEP assembly included the two alleles for 13,183 genes (37.08%) whereas the phased assembly of HiFiAsm recovered two alleles for only 8,484 genes (23,86%). The primary assembly of HiFiAsm, which is expected to be a haploid representation of the genome, only has more than one copy for 156 single copy genes. NGSEP also identified a larger number of multicopy genes compared to HiFiAsm, although the total number of multicopy reconstructed alleles was lower for the NGSEP assembly compared to the HiFiAsm assembly. In terms of computational efficiency, both tools were able to reconstruct the genome in about 50 hours using 32 threads.

**Table 1.** Metrics for diploid assemblies using NGSEP and HiFiAsm over the human HG002 diploid cell line.

| **Metric** | **NGSEP** | **HiFiAsm**<br>**phased** | **HiFiAsm**<br>**primary** |
| --- | --- | --- | --- |
| Length (Mbp) | 5,593.63 | 5,979.17 | 3,109.3 |
| NGA50 (Mbp) | 1.68 | 91.07 | 69.24 |
| Single copy genes in both the reference and the assembly | 15,638 | 23,045 | 34,793 |
| Single copy genes duplicated in the assembly | 13,183 | 8,484 | 156 |
| Single copy genes with exons mapped in different contigs | 209 | 5 | 7 |
| Single copy genes with 50%-99% of sequence mapped | 674 | 93 | 67 |
| Single copy genes with 10% - 50% of sequence mapped | 477 | 23 | 3 |
| Single copy genes with < 10% of sequence mapped | 5,366 | 3,897 | 521 |
| Duplicated genes in the reference found in the assembly | 2,226 | 1,551 | 1,422 |
| Total alleles of duplicated genes | 7,499 | 8,187 | 8,266 |
| Fraction of missing multicopy genes | 0.61 | 0.73 | 0.62 |
| Gene completeness (asmgene) (%) | 81 | 89 | 98 |
| Execution time (h) | 49 | 52 | 52 |

### Benchmark with ONT data

To test the performance of our algorithms with Nanopore reads, we downloaded and assembled datasets of Nanopore reads sequenced from samples of *Escherichia coli, Saccharomyces cerevisiae*, and *Drosophila melanogaster*. We compared the assemblies obtained using Canu, Flye, and NECAT, as well as NGSEP. Figure 5 shows the statistics of these assemblies comparing these tools. Complete assembly statistics are shown in the Supplementary Table T2. For *E. coli*, the most contiguous assembly was obtained with NGSEP after error correction using NECAT. This genome was assembled in one contig by all tools except Flye, which reported two contigs. Using this dataset, the N50 value ranged from 3.57 Mbp (Flye) to 4.62 Mbp (NGSEP). The yeast genome was assembled in its 17 chromosomes by NECAT with an N50 of 0.94 Mbp and by NGSEP with an N50 of 0.81 Mbp after performing error correction with NECAT. The next best tool was Canu, reporting 33 contigs and an N50 of 0.81 Mbp. Finally, Flye assembled the yeast genome in 33 contigs with N50 equal to 0.8 Mbp. The last dataset included in our analyses consisted of reads from the fruit fly. This genome was assembled in 679 contigs (N50 0.91 Mbp) using NGSEP with NECAT error correction, which was the lowest number of contigs obtained. However, Flye achieved an N50 of 1.1 Mbp, being the best result obtained for this dataset. Canu reported 3858 contigs with an N50 of 0.43 Mbp. Unfortunately, NECAT failed to assemble these sequences with the available computational resources, requiring more than 60 GB of RAM memory for this process.

**Figure 5.**
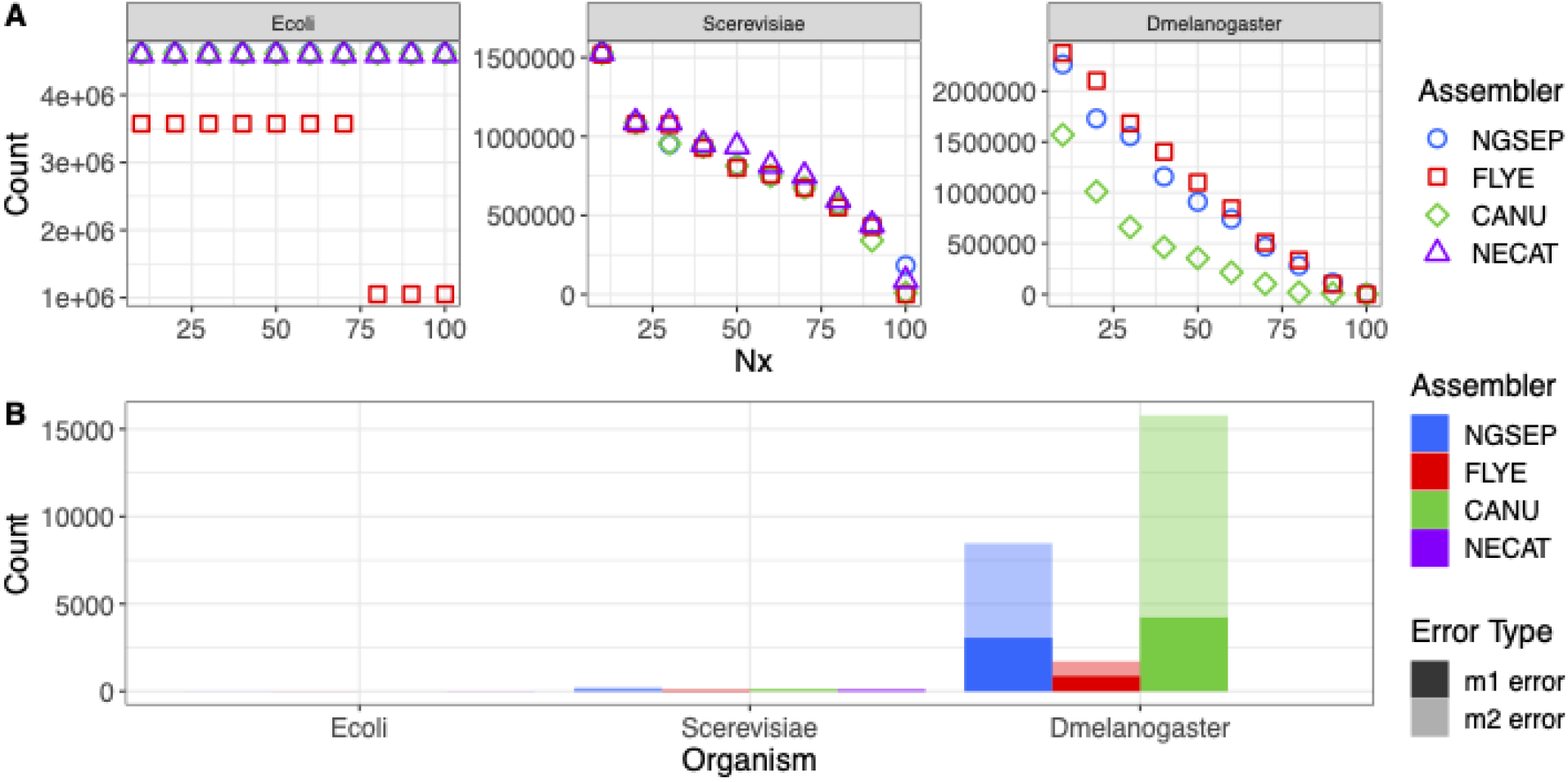
Haploid genomes assembly results using ONT reads. A. Nx curve B. Misassemblies (m1 error) and local misassemblies (m2 error) reported for each genome vs reference genomes.

### Other related features

Based on the development of the genome assembler, version 4 of NGSEP also includes a module to calculate the spectrum of k-mer counts, either from sequencing reads or from a genome assembly. For a k-mer size less or equal to 15, the k-mer counts are stored in a fixed array of 2-byte integers of size 2^30^. This allows to create the spectrum with a fixed RAM usage of 2 gigabytes for an arbitrary number of input reads. Based on this spectrum of k-mers, we included a functionality for error correction in which substitution errors can be corrected by looking at single changes producing k-mers within the distribution of k-mer counts. Moreover, the minimizers table generated to perform efficient identification of read overlaps was also used to create a reference alignment tool for long reads. To keep the algorithm memory tractable, minimizers appearing 1,000 or more times within the reference sequence are discarded. Minimizers for each read are calculated and searched in the minimizers table corresponding to the reference sequence. Minimizer hits are interpreted as k-mer ungapped alignments and clustered according to the read start site predicted for each read. We assessed the performance of the minimizers algorithm implemented in NGSEP for aligning simulated long reads, comparing the results with the alignments obtained using Minimap2 (Li H 2018). Both tools achieved almost perfect accuracy for *S. aureus* and *S. cerevisiae* genomes. Minimap2 showed 3% higher mapping accuracy for the experiment with the human chr20 but NGSEP reported lower root mean squared error (RMSE) values (Supplementary figure S2).

Finally, for circular genomes we implemented a circularization feature as an option of the genome assembler. Given an input set of possible origin sequences, NGSEP maps these sequences to the assembled contigs using the long read alignment algorithm. Each presumably circular contig is rotated and oriented based on the best alignment of an origin sequence.

## DISCUSSION

In this work, we present the results of our latest developments to facilitate de-novo construction of genome assemblies using long reads, which includes novel algorithmic approaches to perform the different steps of the Overlap-Layout-Consensus model. Experiments with a wide variety of datasets indicate that our approach achieves competitive accuracy and efficiency, compared to state-of-the-art tools. From the user perspective, NGSEP achieves nearly perfect assemblies for several species and it is able to reconstruct most gene-rich regions, even in complex genomes. One major advantage of our software is that, combined with previous developments, it offers an easy-to-use, open source and platform independent framework to run a complete analysis of high throughput sequencing reads, including de-novo assembly, read mapping, variants detection, genotyping, and downstream analysis of genomic variation datasets.

The algorithms designed and implemented in NGSEP contribute new alternatives to identify solutions to the genome assembly problem. Although the graph construction with two vertices per read has been used in previous works (Miller et al. 2008, Koren et al. 2017), current software tools seem to implement the classical directed string graph, which requires taking early decisions on the orientation of each read (Cheng et al. 2021). We believe that the undirected graph used in this work makes a better representation for DNA sequences compared to the string graph because it takes into account that DNA is double-stranded and hence it captures more information from the input reads. This allows devising algorithmic approaches different from a greedy traversal of a curated string graph. Moreover, to achieve improved computational efficiency, we avoided complete alignments between reads. Instead, we performed estimations of different types of information (overlap, CSK and percentage of the overlap supported by evidence), that can be used as features to select edges building assembly paths based on a likelihood calculation for each edge. The layout algorithm of NGSEP is inspired by the classical Christofides algorithm for the travel salesman problem, treating the path construction as an edge selection process. Edge features are combined based on their likelihood, replacing edge filtering by edge prioritization. This approach eliminates the need of hard filtering decisions and makes the algorithm adaptable to genomic regions with different repeat structures, as well as to the analysis of reads with variable sequencing error rates.

Taking into consideration Nx curves and misassemblies, NGSEP produces high-quality assemblies with higher contiguity than Flye and a lower number of errors compared to Canu and HiFiASM. These statistics suggest that NGSEP can be used as an accurate alternative to assemble PacBio HiFi reads. Although further work is required to improve N50 in complex assemblies (especially human diploid samples), our results indicate that the contiguity achieved by NGSEP assemblies is enough to reconstruct most gene elements and, moreover, it seems to perform a better allele reconstruction for diploid genomes, compared to HiFiAsm. As shown by recent works (Garg et al. 2021, Nurk et al. 2022, Porubsky et al. 2021), contiguous haploid and diploid assemblies of complex genomes still require the integration of data from technologies or strategies that provide scaffolding and phasing information such as Hi-C or parental sequencing. However, our experiments with diploid samples indicate that new algorithms implemented in existing or novel tools could significantly improve the accuracy of phased assemblies directly from long reads.

Regarding Oxford Nanopore reads with high error rates, NGSEP was able to perform accurate assemblies after reads were corrected running the specialized algorithm implemented in NECAT. This error correction step is crucial in the assembly process of current ONT reads. However, upcoming improvements in the read quality are likely to produce ONT HiFi reads, eliminating the need of a specialized error correction step. We believe that the new algorithms presented in this manuscript make a significant contribution to the development of bioinformatic algorithms and tools for genome assembly. Moreover, the new functionalities of NGSEP facilitate the construction of genome assemblies to researchers working on a wide range of species.

## METHODS

### Benchmark datasets

PacBio and Nanopore publicly available raw datasets were retrieved from NCBI. Haploid datasets included PacBio HiFi/Circular Consensus Sequence (CCS) 20k reads from *Oryza sativa* Indica MH63 accession (<u>PRJNA558396</u>) (Song et al. 2021) and 15k reads from *Oryza sativa* Indica MH63 accession (SRR10188372), PacBio CSS from *Zea mays* B73 accession (<u>PRJNA627939</u>) (Hon et al. 2020), and PacBio CCS from the CHM13 human haploid cell line (<u>PRJNA530776</u>) (Nurk et al. 2022). The human male HG002/NA24385 was used as the diploid dataset (<u>PRJNA586863</u>). Nanopore reads for *Escherichia coli* K12 were obtained from the Loman Lab available at http://lab.loman.net/2015/09/24/first-sqk-map-006-experiment/ (Loman et al. 2015). We selected run MAP-006-1, which also corresponds to the dataset used by Canu in their tutorial. Nanopore reads for *Saccharomyces cerevisiae*, and *Drosophila melanogaster* were directly downloaded from http://www.tgsbioinformatics.com/necat/ (Chen et al. 2021).

### Long read haploid genome assembly tools comparison

We compared the performance of the algorithm described in this work with the algorithms implemented in HiCanu (Nurk et al. 2020), Flye (Kolmogorov et al. 2019), and HiFiASM (Cheng et al. 2021) for PacBio HiFi reads; and with the algorithms implemented in Canu, Flye, and NECAT (Chen et al. 2021) for Nanopore reads. WTDBG (Ruan and Li 2019) was not included because in some initial benchmark experiments it reported a much lower accuracy for complex genomes, compared to other tools, and because it seems to be replaced by HiFiAsm. All PacBio assemblies were run in a Microsoft Azure Standard E64as_v4 (64 vcpus, 512 GiB memory) virtual machine. The parameters used for each tool are detailed in the Supplementary Table T3 and Supplementary Table T4.

### Comparison of genome assemblies with reference genomes

To compare the assembly achieved by each tool against a reference genome, we used Quast with default parameters (Gurevich et al. 2013). Whereas reference coverage, assembly length and N50 were used as sensitivity measures, number and type of misassemblies were used as specificity measures. We calculated and compared these statistics among all assemblies per dataset. The Nx curve was also calculated for each assembly. The reference genomes used in the comparison were *Oryza sativa* Indica MH63 (CP054676–CP054688) (Song et al. 2021), *Zea mays* B73 v.5 (GCA_902167145.1) (Jiao et al. 2017), human haploid line CHM13 v2.0 (https://github.com/marbl/CHM13) (Nurk et al. 2022); the genomes of *Drosophila melanogaster* v.6, *Escherichia coli* K12, and *Saccharomyces cerevisiae* S288c were downloaded from the NECAT web site (Chen et al. 2021).

### Diploid genomes benchmarking

Simulations: To assess the accuracy of the algorithm implemented in NGSEP for reconstruction of diploid samples, we simulated two single chromosome individuals. First, we built a synthetic individual joining two different MHC alleles: the reference allele extracted from GRCh38, and an alternative reconstruction available at the NCBI nucleotide database (accession NT_167249), generated as part of the MHC haplotype project (Horton et al. 2008). Second, we built an individual joining the rice chromosome 9 reconstructions of the reference genome (Nipponbare) and MH63. We simulated 10,000 and 125,000 reads respectively from each simulated diploid individual using the SingleReadsSimulator of NGSEP with average length of 20 Kbp, a standard deviation of 5 Kbp, a substitution error rate of 0.5% and an indel error rate of 1%.

HG002: NGSEP v4.0.1 and HiFiAsm v0.16.0 (Cheng et al. 2021) were employed to obtain a diploid assembly for the Personal Genome Project Ashkenazi Jewish son HG002 (four runs with accession numbers SRR10382244, SRR10382245, SRR10382248 and SRR10382249). We registered time of execution over a node with an AMD EPYC 7402 2.80 Hz, 24C/48T, 128M Cache, a DDR4-3200 processor, 32 cores and 512Gb of RAM. We converted the output files from HiFiAsm (*ctg.gfa) to fasta (*.fa) and merged the haplotypes (*hap1.p_ctg.fa and *hap2.p_ctg.fa) to calculate the main metrics and compare against the NGSEP diploid assembly. Metrics such as N50 and L50 were obtained using Quast v5.0.2. A validation of those metrics was obtained using minigraph v0.19 (Li et al. 2020) and paftools v2.24-r1132-dirty (Li H 2018).

Structural variations are commonly mistaken as misassemblies by current alignment-based evaluations. Hence, the reference-based asmgene method was used to calculate both gene completeness and the number of missing multi-copy genes as additional assembly-quality indicators. According to Cheng et al. 2021, gene completeness equals to |*{SCorMCinASM}*∩*{SCinREF}*|*/*|*{SCinREF}*|, where *{SCinREF}* corresponds to the set of single-copy genes in the reference genome and {SCorMCinASM} refers to the union sets of single-copy and multicopy genes in the assembly. Likewise, missing multi-copy genes are calculated as 1 - |*{MCinASM}* ∩ *{MCinREF}*| / |*{MCinREF}*|. For clarity purposes, a gene is considered as a single copy (SC) if only one match is described into the reference genome (at a 99% of identity), otherwise it is a multi-copy (MC) gene.

### Accuracy assessment for long read alignment

Simulated reads were aligned against their respective reference sequence using Minimap2 v2.17 (Li H 2018) and the ReadsAligner command of NGSEP v4.2.1 with k-mer lengths of 15 (Default mode) and 20. Default parameters were used for all aligners. For time performance evaluation, we conducted all alignments using 4 cores of processing and 20 GB of memory. We evaluated the accuracy of the aligners using percentage of aligned reads, as well as sensitivity and false positive rate metrics. These metrics were calculated using a script that, taking an alignment file as input, infers the real position in the reference genome for each aligned read from the read name and calculates the difference with the position where the read is aligned. Total alignment rate and RMSE are calculated after the total number of aligned reads is counted and the square error rate is totalized over the alignments. This script is available with the NGSEP distribution (class ngsep.benchmark.QualityStatisticsAlignmentSimulatedReads). Accuracy metrics were computed for bam files filtered by alignment quality values from 0 to 80.

## ACKNOWLEDGEMENTS

This work has been supported by the Colombian research fund “PATRIMONIO AUTÓNOMO FONDO NACIONAL DE FINANCIAMIENTO PARA LA CIENCIA, LA TECNOLOGÍA Y LA INNOVACIÓN FRANCISCO JOSÉ DE CALDAS” through the grant with contract number 80740-441-2020, awarded to JD. This work was also supported by internal funds of Universidad de Los Andes through the FAPA initiative led by the Vice-presidency of Research and Knowledge Creation. We also wish to acknowledge the support of the IT Services Department and ExaCore-IT Core-facility of the Vice Presidency for Research & Creation at the Universidad de Los Andes that allow us to perform the computational analysis.

